# Glycolysis and interferon transcription in melanoma immunotherapy

**DOI:** 10.64898/2026.10.30.763008

**Authors:** Lili Wu, Wei Yang

## Abstract

Glycolysis and interferon signalling have both been associated with responses to PD-1 blockade, but their joint transcriptional relationship with treatment outcome remains uncertain. We examined whether a state with high glycolysis and low interferon-α response was associated with reduced disease control in melanoma. Fixed Hallmark gene sets were scored by single-sample gene set enrichment analysis in GSE91061, GSE78220 and an additional Liu cohort. Disease control analyses included 47, 26 and 100 patients. Conditional odds ratios for the joint state were 6.06 (exact 95% confidence interval, 1.04 to 65.61), 0.35 (0.026 to 3.11) and 1.03 (0.39 to 2.73), respectively. Holm-adjusted P values for the two initial comparisons were 0.073 and 0.365. The additional exploratory P value was 1.000. Continuous models and analyses that omitted each patient in turn did not provide consistent evidence of an adverse association. IFNα scores were negatively correlated with DNA-derived tumour purity in 102 patients (Spearman ρ = −0.671). Among 40 patients with paired biopsies, disease control was associated with higher on-treatment IFNα scores after baseline adjustment, although the paired subset had lower baseline glycolysis than patients without paired profiles. In patient-level single-cell analyses, the baseline CD8 T cell correlation weakened when an alternative rank score used a common gene background. The defined joint state was not consistently associated with poorer disease control across cohorts. Bulk IFNα scores were related to tumour purity, and cellular correlations were sensitive to scoring choices. These transcriptomic results do not measure pyruvate concentration, glycolytic flux or STAT1 pyruvylation.

## Introduction

PD-1 blockade produces durable responses in some patients with melanoma, whereas others progress during treatment. Tumour metabolism has been investigated as one source of this variation. Harel and colleagues linked metabolic proteins with antigen presentation and immunotherapy outcomes [1], and Mahmood and colleagues found that mitochondrial DNA perturbations increased glycolytic flux and enhanced checkpoint blockade response in melanoma models [2]. Metabolic expression states have also been used to study treatment response in immune cells [3]. Bulk tumour RNA, however, combines malignant, immune and stromal expression. Single-cell studies have identified distinct tumour and T cell programmes associated with checkpoint blockade outcomes [4,5]. A metabolic score calculated from bulk RNA can therefore reflect more than the metabolic state of malignant cells.

Zuo and colleagues reported that glycolysis associated with high glucose promoted STAT1 pyruvylation at Lys201, impaired the interaction between STAT1 and STAT2 and suppressed type I interferon-mediated antiviral activity [6]. This finding suggests that high glycolysis transcription combined with low interferon response transcription might identify an adverse state in melanoma immunotherapy. The proposed relationship requires direct examination in clinical data. The pyruvylation mechanism was studied in settings involving high glucose and antiviral responses, and an RNA score cannot quantify the relevant protein modification. Moreover, prolonged tumour interferon signalling can promote resistance to checkpoint blockade [7]. The direction of a clinical association cannot be inferred from the biochemical mechanism alone.

The timing of sampling and the cell population being measured may also affect the comparison. On-treatment pathway signatures have previously been associated with anti-PD-1 response in melanoma [8]. Baseline and on-treatment expression consequently address different clinical questions. Differences between glycolysis and interferon scores may additionally arise from gene set overlap, the genes available for ranking and the numbers of cells contributing to an expression aggregate. These factors are particularly relevant when small patient groups are compared across bulk and single-cell datasets.

In this study, we examined whether a transcriptional state with high glycolysis and low IFNα response was associated with disease control in GSE91061 and GSE78220, followed by an additional evaluation in the Liu cohort [9,10,11]. The two programmes were defined by fixed Hallmark gene sets and scored with ssGSEA [12,13,14]. We compared median-based states with continuous models and assessed the influence of individual patients, tumour purity and paired-biopsy changes. We then analysed GSE115978 and GSE120575 at the patient level to compare malignant and immune cell expression [4,5]. Alternative rank scores and gene backgrounds were used to examine the sensitivity of the correlations between glycolysis and IFNα.

## Materials and methods

### Study design and analysis definitions

The present computational study was based on three published bulk melanoma cohorts and two single-cell resources. The initial primary comparison was disease control in the state with high glycolysis and low IFNα response versus the other three states, separately in GSE91061 and GSE78220. We documented the initial definitions retrospectively and recorded subsequent analysis amendments. Continuous interactions, covariate adjustment, paired-biopsy analyses and single-cell comparisons were exploratory. We retained the original primary tests and their two-test multiplicity family. The Liu evaluation and further sensitivity analyses were added after the initial two-cohort results were available. Its four inclusion populations, endpoints and thresholds were fixed before associations between scores and outcomes were calculated. These amendments are identified in the supporting protocol. No gene set, threshold, endpoint or population was selected by association strength. The study assessed associations rather than training a clinical prediction model.

### Expression data and clinical endpoints

GSE91061 contained 109 RNA sequencing profiles from 65 patients treated with nivolumab, including 51 baseline profiles [9]. Its clinical table contained additional patients without corresponding RNA profiles and was not used to define the number of patients in the expression cohort. GSE78220 supplied 28 expression records [10]. We removed the early on-treatment Pt16.OnTx record and averaged the two baseline Pt27 biopsies on the linear FPKM scale, leaving 26 baseline patients. We matched the expression and clinical records by patient identifier. In the source processing, GSE91061 Entrez identifiers were mapped to gene symbols, and 121 unmatched identifiers were dropped. GSE78220 symbols were mapped using official symbols and unambiguous aliases. Ambiguous and unmatched mappings were excluded. Duplicate symbol rows were averaged on the linear scale. The resulting matrices contained 21,840 and 23,575 rows. Mapping tables are retained in the accompanying data package.

Disease control was defined as complete response, partial response or stable disease, with progressive disease coded as no disease control. Objective response was restricted to complete or partial response. Responder labels assigned by the original authors were not assumed to be interchangeable with these endpoints. Ambiguous or conflicting entries for best response were excluded from the affected endpoint. Four baseline GSE91061 patients did not enter the primary analysis. GSE91061 used RECIST v1.1 and GSE78220 used immune-related RECIST. Harmonizing response categories did not make these assessment criteria identical. Previous exposure to MAPK inhibitors in GSE78220 was verified against GEO sample characteristics. There were 11 patients with previous exposure and 15 without it. These variables were used in a separate exploratory adjustment, without treating multiple anatomical sites from one patient as independent observations.

The Liu cohort [11] supplied a released TPM matrix for 121 patients and 20,848 unique genes, matched to Supplemental Table 1 by patient identifier. The source matrix had already been restricted to genes expressed in at least 25% of tumours. We transposed it without changing TPM values. All patients received nivolumab or pembrolizumab. The primary added population required source code 3 for biopsy context, a biopsy on or before PD-1 initiation and available expression. We conservatively excluded Patient143, Patient107 and Patient61 because the text describing RNA quality control named them as excluded, although they remained in the released matrices. This yielded 102 patients for thresholds and 100 for disease control and objective response after exclusion of mixed responses. The three sensitivities were context 3 using all released profiles (104/102 threshold/endpoint patients), all pre-PD1 biopsies excluding the three quality conflicts (117/115), and all pre-PD1 released profiles (120/118). Patient56, biopsied five days after therapy began, was excluded from every baseline population. Mixed responses contributed to baseline thresholds and standardization but not endpoint models. All four scenarios were reported. The Liu and Riaz data were released by separate studies, and Liu and colleagues used the Riaz cohort for external validation. Patient identifiers could not be matched across the resources, so individual overlap could not be excluded. In the conservative population designated by the source as pre-PD1, biopsies ranged from 490 days before therapy to the day of initiation. This source category does not imply a uniform or recent sampling interval.

**Table 1.** Datasets and statistical units included in this reanalysis.

| Dataset | Source material | Analysis population | Role |
|---|---|---|---|
| GSE91061 | 109 bulk RNA profiles / 65 patients | 51 at baseline, 47 with resolved response, 40 paired with response | Primary comparison and paired analysis |
| GSE78220 | 28 bulk expression records | 26 baseline patients after visit exclusion and biopsy averaging | Second cohort comparison |
| Liu 2019 | 121 released bulk RNA profiles | 102 in the conservative baseline population, 100 with resolved response | Additional external evaluation |
| GSE115978 | 7,186 annotated cells | Verified donor mapping. Untreated malignant 9 / CD8 10 / myeloid 4 patients | Exploratory cellular analysis by patient |
| GSE120575 | 16,291 immune cells | Anti-PD-1 at baseline, 12 CD8 / 6 myeloid patients. During treatment, 19 / 11 | Immune cell analysis by patient |

Counts for different target types may overlap within patients and must not be summed. Cell totals do not represent the number of independent patients. External cohort eligibility and the discrepancy in quality exclusions between the publication and matrix are detailed in Supplementary Table S1. GSE120575 contains sorted immune cells, so cell counts do not estimate tissue cell fractions.

### Calculation of transcriptional scores

The human MSigDB Hallmark collection, release 2026.1.Hs, was used to define the transcriptional programmes [12]. The two main gene sets were HALLMARK_GLYCOLYSIS and HALLMARK_INTERFERON_ALPHA_RESPONSE. IFNγ response, hypoxia, E2F targets and G2M checkpoint were retained for contextual analyses. Single-sample GSEA was implemented with GSEApy [13,14], using rank sample normalization, rank correlation normalization, weight 0.25, gene set size limits of 15 to 500, the full available gene background and no permutation-based enrichment testing. All analyses used the enrichment score (ES). Normalized enrichment scores were not used. The original records used GSEApy 1.1.9. An independent bulk rescore with version 1.3.1 reproduced all 462 reported scores to less than 10^−9^ absolute difference. Measured gene coverage was 199/200 and 200/200 for glycolysis and 95/97 and 97/97 for IFNα in GSE91061 and GSE78220, respectively. All on-treatment samples were excluded when defining baseline thresholds.

Within each cohort, a score strictly above its baseline median was classified as high. Ties were classified as low. The glycolysis and IFNα thresholds were 5,078.584 and 6,342.876 in GSE91061, and the corresponding thresholds for GSE78220 are provided in the source data. Medians used all patients scored at baseline, including those later excluded from the endpoint comparison. Thus, the classifier is defined relative to each cohort and was not evaluated as a fixed threshold transferable to new patients. Calculations used the original ES values.

For the Liu matrix, the same NCBI gene information reference and rules for official symbols and unambiguous aliases used in GSE78220 mapped 19,889 rows directly and 927 through aliases. We dropped 29 ambiguous and three unmatched rows and averaged 14 duplicate mapped rows on the linear TPM scale, leaving 20,802 genes. The mapping decisions and reference checksum are provided with the analysis records. The mapping implementation exactly reproduced the earlier GSE78220 mapping table. External coverage was 200/200 glycolysis and 96/97 IFNα genes. No additional expression filter or log transformation was applied. Each external scenario used all its eligible baseline patients, including mixed responses, for medians and standardization. Thresholds remained outcome independent.

The initial mapping contained one collapsed nonsymbol row labelled “-” in each bulk matrix. A sensitivity analysis removed this row and repeated scoring and median classification. No baseline patient changed joint state assignment in either cohort, and both primary exact tables and estimates were identical. Independent rescoring also reproduced all 258 glycolysis, IFNα and IFNγ values from paired visits across 43 complete patient pairs to less than 10^−9^ absolute difference. The sensitivity analysis of input mapping and the complete score comparisons accompany the scripts.

### Statistical comparisons and regression analysis

For each 2 × 2 table, we used a two-sided Fisher exact test and a conditional maximum likelihood odds ratio with an exact conditional 95% confidence interval. The odds ratio compares the odds of disease control in the joint state group with those in the other states. Holm adjustment covered the two primary P values from the cohort comparisons. The confidence intervals are individual, unadjusted intervals. Their exclusion of one need not agree with a multiplicity-adjusted P value. Two-sided tests were used throughout. A threshold of 0.05 was used for the corrected primary tests, without interpreting nonsignificance as evidence of no effect.

The original exploratory logistic models of main effects and models adjusted for one covariate used ordinary maximum likelihood and Wald intervals. Continuous scores were standardized using the mean and sample SD of all eligible baseline patients in each cohort or external inclusion scenario. The original models adjusted for one covariate included the joint state and one of the immune score, hypoxia, E2F, G2M, IFNγ or prior treatment relevant to the cohort. The 12 joint state effects formed a separate Benjamini-Hochberg family. Added continuous models included both standardized scores and their product. Firth penalized logistic likelihood and penalized profile intervals addressed sparse data [15,16]. A chi-square reference supplied approximate likelihood ratio tests. Holm correction covered the continuous interactions in the two original cohorts and the interaction between cohort and state as a separate family of three tests. The cohort interaction estimates a ratio of state odds ratios, not a pooled effect. An exact conditional analysis fixed each cohort’s table margins and total joint state successes and enumerated the remaining support, with a probability-ordered two-sided P value and central exact interval [17]. Implementation checks included saturated two-by-two closed-form estimates, analytic-score finite differences, profile endpoints and optimizer convergence. Ordinary maximum likelihood estimates were retained as sensitivity analyses. Each patient was also omitted once, with thresholds and standardization held fixed. These fits assess influence, not predictive performance.

The added external analysis reports conditional exact odds ratios and two-sided Fisher tests for disease control and objective response under every predefined eligibility scheme. These nominal P values are exploratory, with no retroactive change to the original primary family. Firth models of main effects and continuous score products used standardization over all baseline patients in each scenario. Ordinary maximum likelihood fits were retained as sensitivities. In the conservative external population, three predefined models included the joint state and adjusted for prior CTLA-4 blockade, DNA-derived purity or both. Purity was standardized over all 102 eligible baseline patients. All 100 patients with evaluable endpoints had both covariates, so no complete cases were lost. All model coefficients, diagnostics and nominal tests are disclosed. No added result is used as confirmatory evidence.

### Immune scores and tumour purity

ESTIMATE immune and stromal scores [18] were supplied by the tidyestimate implementation in the analysis package. We independently joined these scores to the baseline patient table and recalculated Spearman correlations with the two main ES values. We did not transform these scores into calibrated estimates of tumour purity from RNA sequencing. The immune and pathway scores were derived from the same expression input. Their correlations therefore describe relationships between transcriptional summaries and do not provide independent measurements of cell fractions. IFNα/IFNγ overlap was calculated among measured genes in each cohort. This analysis did not establish cytokine-specific pathway activation.

The Liu clinical table supplied DNA-derived ABSOLUTE tumour purity. The source article describes purity and ploidy estimation from variant allele frequencies and somatic copy number alterations [11,19]. We calculated Spearman correlations in complete cases between purity and the two main RNA scores in the conservative baseline population and applied Holm correction to these two exploratory tests. Purity was considered an independent DNA-derived admixture estimate, not a direct measure of immune cell abundance or a proof of a causal composition effect. The external adjustment model jointly included the binary state, prior CTLA-4 blockade and purity. Models adjusted for one covariate were retained. Covariates and the handling of complete cases were fixed before inspecting the added associations.

### Analysis of paired biopsies

For the longitudinal comparison, baseline and on-treatment GSE91061 profiles were matched within each patient. Forty-three patients had both visits and 40 had unambiguous disease control labels. Changes were defined as on-treatment minus baseline ES. Means and mean differences were accompanied by percentile bootstrap confidence intervals from 100,000 patient resamples. Group contrasts resampled patients independently within outcome groups. All random seeds are retained in the analysis code and result table. We fitted ordinary least squares to on-treatment IFNα ES with baseline IFNα ES, baseline glycolysis ES and disease control status as covariates, using HC3 heteroscedasticity-robust standard errors and 95% confidence intervals based on a normal reference distribution. No model term was removed on the basis of its P value. Continuous coefficient rescaling for display is specified in the figure legend. This is an exploratory conditional association model, not a model for prospective treatment response.

To assess selection into the longitudinal analysis, we described all 43 patients with paired profiles versus the eight without them, and the 40 paired patients with resolved outcomes versus the seven patients with evaluable outcomes without an on-treatment profile. Continuous standardized mean differences divided the difference in means by the square root of the average of the two group variances. We reported frequencies of prior treatment and outcomes without using significance tests to declare balance. Exact biopsy intervals were not established for every patient. No uniform duration of treatment exposure was assumed.

### Single-cell processing and patient aggregation

The published processed expression matrices and author annotations were used for the single-cell analyses [4,5]. GSE115978 log_2_(TPM/10 + 1) and GSE120575 log_2_(TPM + 1) values were inverted to linear TPM. Additional checks required matched metadata, no invalid expression and at least 500 detected genes per cell. All 23,477 deposited cells passed. Quality control of the sequencing data was not rerun. Eligibility was determined from the number of unique cells in each specimen and target population. Expression was averaged within each specimen and target type, and groups with at least 20 cells were scored. Thresholds of 10 and 50 were explicit sensitivities. These normalized expression aggregates are not count-based pseudobulk measurements. The dataset-specific backgrounds contained 23,686 and 55,737 genes. Absolute scores were not compared across these backgrounds.

In GSE120575, CD8 T cells followed the explicit author Table S2 cell list and myeloid cells pooled annotated monocyte and dendritic cell groups. Cells lacking a supported target annotation were excluded from target analyses. Scores from multiple sorts were averaged equally within lesion, followed by equal lesion averaging within patient, visit and regimen. AntiPD1 monotherapy was separated from combination treatment. Each patient contributed once within a target/visit analysis. The recovered original GSE115978 Table S1A identified repeated material. Mel129PA/PB shared patient Mel129 and Mel75/Mel75.1 shared patient Mel75. The Mel75 materials were separated using the source cohort labels assigned to each cell, reconstructed and scored separately before equal patient averaging. Treatment-naive and previously treated observations were kept in separate strata defined by the source. Heterogeneous prior treatments and incomplete donor mapping across resources prevent treating the two resources as independent efficacy validations.

The retained ssGSEA calculations used lexicographic gene order for deterministic ties. To test robustness independently, we ranked each aggregate over its gene background using average ranks for ties, defined each midpoint percentile as (rank − 0.5) divided by background size, and averaged those percentiles over measured genes in each fixed set. This score requires no arbitrary ordering among equal values. We repeated it with each dataset’s full background and the intersection of 21,957 measured genes. No zero-filled union was used. Gene-order permutations verified numerical invariance. Both backgrounds, every cell threshold and sensitivity analysis of donor mapping were retained. The common background changes the rank reference and some set coverage, including 95 versus 94 measured IFNα genes in GSE120575. It was not assumed to be uniquely correct.

### Single-cell uncertainty and sensitivity analysis

Spearman estimates calculated at the patient level were accompanied by the original asymptotic two-sided P values. The original two baseline GSE120575 target tests retained their Benjamini-Hochberg family. Additional analyses used exact pair permutations when computationally feasible and otherwise 19,999 Monte Carlo pair permutations with a plus-one correction, and 5,000 patient bootstrap resamples for percentile intervals. Seeds, valid bootstrap counts and inference status are retained in the output tables. Strata with fewer than eight patients were descriptive, including the baseline myeloid group of six patients. The supporting tables distinguish asymptotic and permutation P values and report both correction within each stratum and a broad correction across all added tests at the patient level. The sensitivity analyses used overlapping observations and were not regarded as independent replications. We did not compare separate P values to infer a difference between cell types, methods or time points, and the small number of paired patients did not support a treatment-induced correlation claim.

### Reproducibility and use of computational assistance

Processed expression and clinical data are available from GEO (GSE91061, GSE78220, GSE115978 and GSE120575) and the Liu article’s Supplementary Data 2 and Supplementary Tables [11]. The accompanying package provides source checksums, clinical mapping, amendment protocols, patient-level data and scripts for processed-data reconstruction, analysis and figures. The workflow began with processed expression data and did not reprocess sequencing reads. Survival was excluded because the time origins of the available follow-up fields could not be independently verified. Diagnostic records are retained in the analysis package.

The recorded analysis environment used Python 3.12.14, NumPy 2.5.4, pandas 3.0.6, SciPy 1.18.1, statsmodels 0.15.0 and GSEApy 1.3.1. Figures were generated with Matplotlib 3.11.2. Analysis-specific software versions, random seeds and the previous computational verification records are included in the code package. OpenAI ChatGPT/Codex assisted with code drafting, literature checks and manuscript preparation. Procedural guidance from Scientific Agent Skills was also used [20]. Responsibility for the study, interpretation and submitted text remains with the authors.

## Results

### Comparison of the joint transcriptional state across clinical cohorts

GSE91061 provided 51 baseline patients, of whom 47 had an unambiguous disease control endpoint. GSE78220 provided 26 baseline patients after removal of an early on-treatment record and averaging of two baseline biopsies from one patient. The median thresholds were calculated before endpoint exclusions (Table 1).

The group with high glycolysis and low IFNα response contained 13 of 51 and 6 of 26 baseline patients, with 11 and 6 entering the corresponding disease control comparisons (Fig 1a,b).

**Fig 1.**
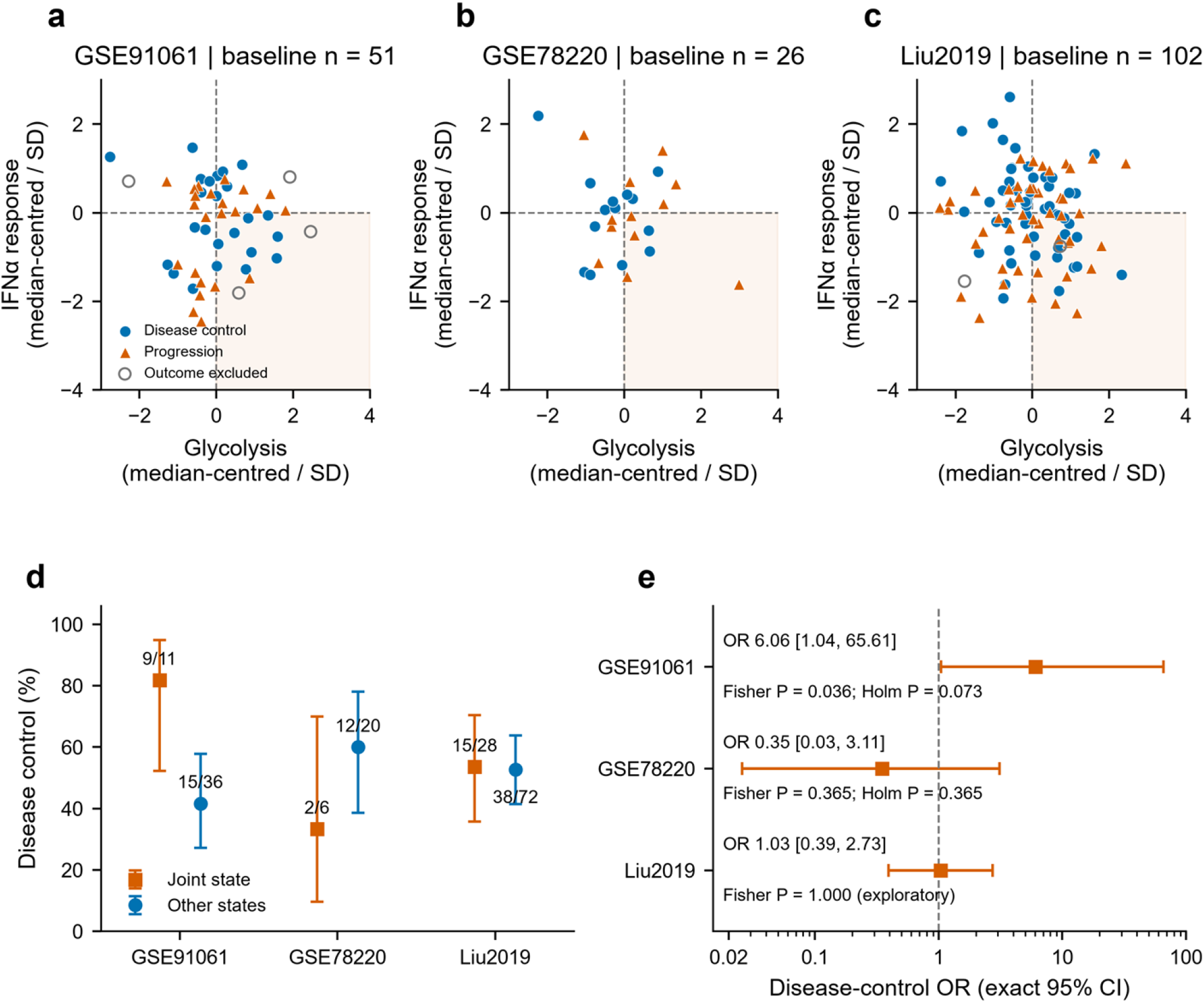
Comparison of the joint state and disease control across cohorts. a to c, Baseline glycolysis and IFNα response ssGSEA scores in GSE91061 (51 patients), GSE78220 (26) and the conservative external Liu population (102). Each score is centred on the cohort/scenario median and divided by its baseline sample SD. Dashed lines are classification thresholds. The shaded lower-right quadrant denotes glycolysis high with IFNα low. Blue circles indicate disease control, orange triangles progression and open circles unresolved or mixed outcomes excluded from the endpoint. d, Disease control proportions and Wilson 95% intervals. Labels show successful outcomes/eligible patients. e, Conditional odds ratios and exact conditional 95% intervals. OR > 1 denotes higher disease control odds in the joint state. The original two cohorts show raw Fisher and original two-test Holm P values. The Liu comparison shows a separately labelled exploratory raw P value. Eligibility rules for the Liu evaluation were fixed before associations between its scores and outcomes were inspected. Each patient enters once per cohort. No pooled effect is shown.

In GSE91061, disease control was observed in 9 of 11 patients in the joint state group and 15 of 36 patients in the other states. The conditional odds ratio was 6.06 (exact 95% confidence interval, 1.04 to 65.61, Fisher P = 0.036). Its direction was opposite to the proposed adverse association. In GSE78220, the corresponding counts were 2 of 6 and 12 of 20, giving an odds ratio of 0.35 (0.026 to 3.11, P = 0.365). After Holm adjustment for the two initial comparisons, the P values were 0.073 and 0.365. Thus, the initial cohorts did not show a consistent association, and neither comparison met the corrected 0.05 threshold (Table 2, Fig 1d,e).

**Table 2.** Initial and added disease control comparisons.

| **Dataset** | **Joint state<br>DC / n** | **Other states<br>DC / n** | **Conditional OR (95% CI)** | **Raw P / original Holm P** |
|---|---|---|---|---|
| GSE91061 | 9 / 11 | 15 / 36 | 6.06 (1.04 to 65.61) | 0.036 / 0.073 |
| GSE78220 | 2 / 6 | 12 / 20 | 0.35 (0.026 to 3.11) | 0.365 / 0.365 |
| Liu 2019 | 15 / 28 | 38 / 72 | 1.03 (0.39 to 2.73) | 1.000 / not in original family |

The joint state denotes high glycolysis and low IFNα response relative to the baseline medians in each cohort. DC denotes complete response, partial response or stable disease. The comparator outcome is progressive disease. Odds ratios and confidence intervals are conditional exact estimates. P values are two-sided Fisher tests. Holm adjustment covers the two cohort tests, while the displayed 95% intervals are not multiplicity-adjusted. The Liu comparison was added after the initial analysis with rules fixed before its associations were inspected. Its nominal P value is exploratory and it is not retroactively included in the original two-test family. All four external eligibility scenarios are retained in the supporting information.

The Liu cohort was examined as an additional external evaluation with eligibility rules fixed before associations between its scores and outcomes were calculated. Of 121 matched patients, the conservative baseline population included 102 after applying the source biopsy category and excluding reported failures of RNA quality control. Two records with mixed responses contributed to the thresholds but were excluded from the endpoints, leaving 100 patients. Disease control occurred in 15 of 28 joint state patients and 38 of 72 other patients. The conditional odds ratio was 1.03 (exact 95% confidence interval, 0.39 to 2.73, exploratory Fisher P = 1.000, Fig 1c to e).

The source publication and released matrix disagreed about three quality exclusions. To examine this uncertainty, all four defined eligibility scenarios were retained. The three additional scenarios contained 102, 115 and 118 patients with evaluable endpoints. Across the four scenarios, disease control odds ratios ranged from 0.92 to 1.04, with wide intervals. These are overlapping sensitivity populations, and their results do not establish equivalence or provide additional independent replications (Supplementary Table S1).

The distribution of response categories differed between the cohorts. GSE91061 contained 14 cases of stable disease, whereas GSE78220 contained none. Disease control and objective response were consequently identical in GSE78220. Odds ratios for objective response were 2.78 (0.45 to 16.04), 0.35 (0.026 to 3.11) and 1.76 (0.66 to 4.69) in the three cohorts. Two GSE91061 patients had stable disease in the BOR field but progression in IBOR. Giving either field priority preserved the direction of the joint state association while changing its uncertainty. Both alternatives are reported without replacing the initial primary population (Supplementary Tables S1 and S2, S1 Fig).

### Continuous models and the influence of individual patients

To examine whether the categorical result depended on the median classification, we analysed standardized glycolysis and IFNα scores together with their product. Firth interaction odds ratios were 0.70 (95% penalized profile interval, 0.28 to 1.38) in GSE91061 and 1.00 (0.56 to 2.23) in GSE78220. The corresponding additional Liu estimate was 0.71 (0.45 to 1.04, exploratory penalized likelihood P = 0.080). All three interaction intervals included one, providing no consistent evidence that the product of the continuous scores was associated with disease control (Supplementary Table S3).

The difference between the two original cohort-specific associations was assessed using an interaction between cohort and state. A Firth model containing cohort, joint state and their product gave a ratio of the GSE91061 odds ratio to the GSE78220 odds ratio of 13.95 (1.45 to 172.27). The raw penalized likelihood P value was 0.022, whereas the Holm P value across the three added interaction tests was 0.067. Exact conditional enumeration gave a ratio of odds ratios of 15.23 (exact 95% interval, 0.85 to 409.30, P = 0.064). Although the point estimates had opposite directions, the wide intervals and method-dependent inference limited the evidence for a difference between cohorts (S2 Fig).

Each patient was removed once to assess the influence of individual observations. With the original thresholds held fixed, conditional odds ratios ranged from 5.40 to 11.99 in GSE91061 and from 0.18 to 0.46 in GSE78220. Neither direction reversed. However, raw Fisher P values in GSE91061 ranged from 0.011 to 0.071, indicating that nominal significance depended on individual patients. The continuous interaction close to zero in GSE78220 changed sign in 12 of 26 deletion fits. These comparisons assess influence and do not estimate cross-validated prediction performance (Supplementary Table S4, S3 Fig).

The joint state estimates were also examined after covariate adjustment. Separate models adjusted for one covariate in the initial cohorts retained opposite joint state directions, with ordinary maximum likelihood odds ratios of 6.30 to 10.91 and 0.24 to 0.46. None of the 12 state effects met a separate Benjamini-Hochberg threshold of 0.05, and these individual adjustments did not jointly control confounding. In the conservative Liu population, the fixed Firth model including prior CTLA-4 blockade and DNA-derived tumour purity used all 100 patients with evaluable endpoints and gave a state odds ratio of 1.78 (0.69 to 4.74). The adjusted Liu interval remained wide and included one. Results from every specified adjustment are reported in Supplementary Table S3.

### Relationships with immune transcription and tumour purity

We examined immune scores and DNA-derived tumour purity to help interpret the bulk RNA results. IFNα response scores correlated positively with ESTIMATE immune scores in GSE91061 (n = 51, Spearman ρ = 0.742, P = 4.76 × 10^−10^) and GSE78220 (n = 26, ρ = 0.791, P = 1.47 × 10^−6^). Glycolysis correlations with the same immune scores were −0.021 and −0.100 (Fig 2a,b,d,e). Because the scores used the same RNA input, their correlation alone could not identify an effect of cellular composition.

**Fig 2.**
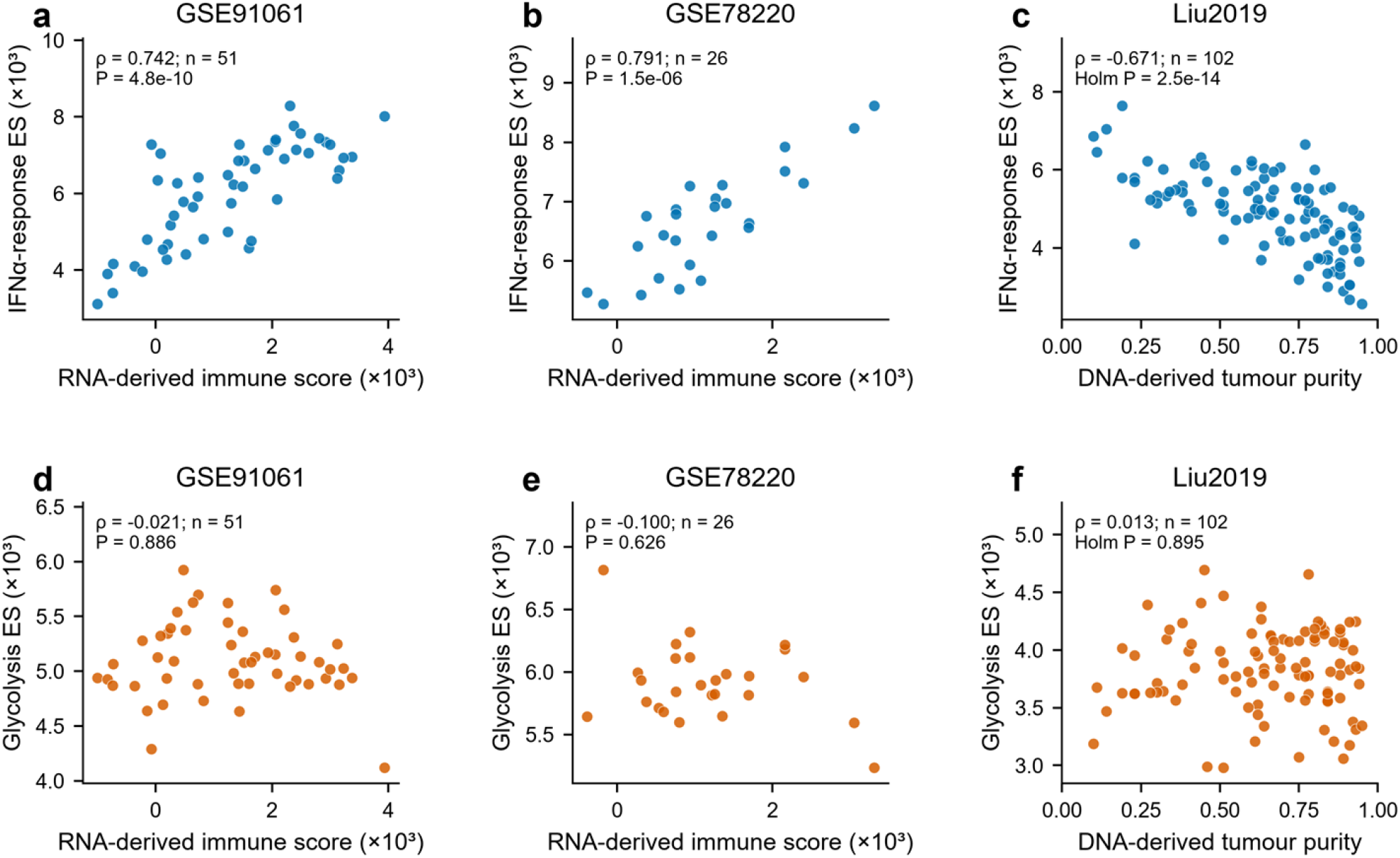
Relationships of transcriptional scores with immune scores and tumour purity. a,b,d,e, Associations of IFNα response and glycolysis scores with an RNA-derived ESTIMATE immune score in GSE91061 (51 patients) and GSE78220 (26). c,f, The corresponding scores against source-reported DNA-derived ABSOLUTE tumour purity in the conservative external Liu population (102 patients). Each point is one baseline patient. ES and immune scores are divided by 1,000 for display only. Purity is the source-reported fraction. Annotations show Spearman correlations and two-sided P values. The two external purity tests show Holm-adjusted P values. RNA-derived scores share their input, whereas the external purity estimate uses independent DNA information. Neither comparison directly measures immune cell fractions or establishes a causal composition effect. Absolute ES values should not be compared across cohort-specific gene backgrounds.

The Liu cohort provided ABSOLUTE tumour purity estimated from DNA alterations in the source study. Among 102 baseline patients, purity correlated negatively with IFNα scores (ρ = −0.671, patient bootstrap 95% interval, −0.768 to −0.542, two-test Holm P = 2.5 × 10^−14^). Its correlation with glycolysis was close to zero (ρ = 0.013, Holm P = 0.895, Fig 2c,f). Tumour purity, however, is not a direct measurement of immune cell fraction. The association and adjusted clinical model do not show that admixture caused the different outcome estimates across cohorts.

The two interferon gene sets also overlapped substantially. The measured IFNα and IFNγ Hallmark sets shared 71 genes in GSE91061 and 73 in GSE78220, leaving 24 measured IFNα-specific genes in each cohort. The IFNα response score consequently cannot be regarded as a cytokine-specific assay. Gene set overlap limits attribution to a specific interferon, while mixed cellular expression limits attribution to a specific cell population.

### Comparison of interferon transcription in paired biopsies

Paired biopsies were used to examine whether IFNα transcription differed over the course of treatment. GSE91061 contained 43 patients with baseline and on-treatment profiles. Forty had resolved disease control labels, including 22 with disease control and 18 with progression. Ten of these 40 patients belonged to the baseline joint state group. The mean change in IFNα score was 426 enrichment score units (patient bootstrap 95% confidence interval, 83 to 793). Mean changes were 717 (264 to 1,239) in patients with disease control and 71 (−406 to 546) in patients with progression. The difference in mean change between the groups was 646 units, with an interval that included zero (−19 to 1,344, Fig 3a to c).

**Fig 3.**
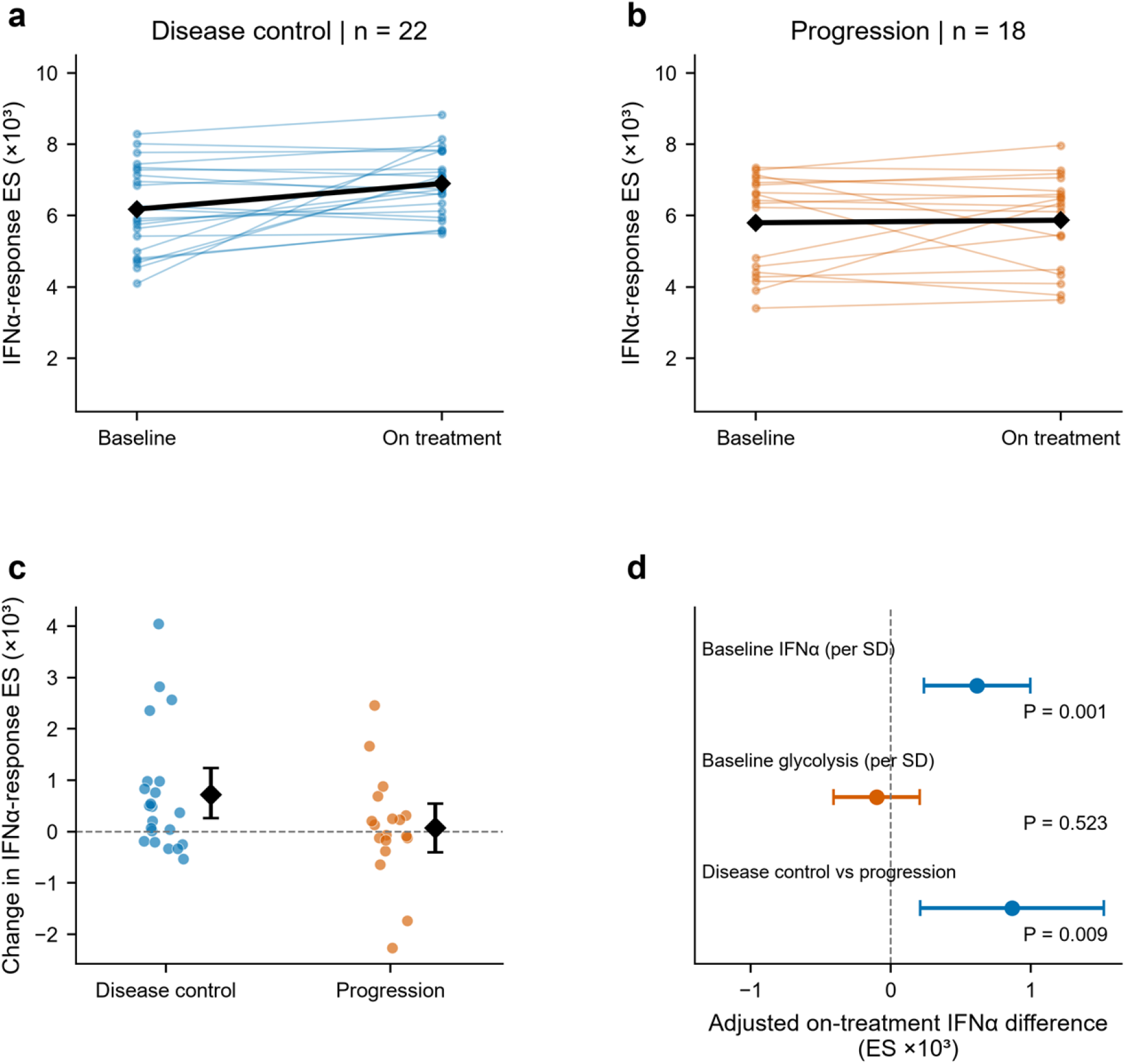
Paired interferon transcription and disease control in GSE91061. a,b, Paired baseline and on-treatment IFNα response ES for 22 patients with disease control and 18 with progression. Coloured lines connect individual patients. Black diamonds and lines show group means. The same y-axis limits are used. c, Individual changes (on treatment minus baseline) with group means (black diamonds) and percentile 95% intervals from 100,000 patient bootstraps. Horizontal jitter is deterministic and used only for visibility. d, Coefficients from the n = 40 ordinary least squares model of on-treatment IFNα ES on baseline IFNα, baseline glycolysis and disease control status. Bars are HC3 robust 95% intervals. Continuous covariate coefficients are displayed per sample standard deviation of the corresponding baseline score among these 40 patients. All ES axes use units of 10^3^. P values are exploratory and unadjusted. This model estimates an on-treatment association, not prospective discrimination. Availability of paired biopsies was associated with a different baseline glycolysis distribution. Supplementary Table S6 quantifies this difference between the selected subset and the remaining patients.

On-treatment IFNα transcription was also analysed with adjustment for baseline IFNα and glycolysis. In this model, disease control was associated with an adjusted difference of 867 score units (HC3 robust 95% confidence interval, 212 to 1,522, P = 0.0095). Baseline IFNα was associated with the on-treatment score, with a coefficient of 0.476 per score unit (0.182 to 0.770, P = 0.0015). The baseline glycolysis coefficient was −0.335 (−1.364 to 0.694, P = 0.523, Fig 3d). The change comparison measured the difference between visits. The adjusted model compared on-treatment scores among patients with similar baseline scores. Neither analysis established prospective prediction or mediation.

The 40 patients with paired biopsies had lower baseline glycolysis than the seven patients with evaluable outcomes without an on-treatment profile (standardized mean difference, −0.97). Prior ipilimumab and disease control each occurred in 22 of 40 paired patients and 2 of 7 unpaired patients. The comparison of all 43 paired patients with the eight unpaired baseline patients gave a glycolysis standardized difference of −1.04 (Supplementary Table S6). The longitudinal result is therefore an exploratory association in a selected subset. It cannot establish a treatment-induced pathway from baseline metabolism through interferon to clinical outcome.

### Single-cell correlations at the patient level and scoring sensitivity

Single-cell data were examined to determine whether the transcriptional relationships could be assigned to particular cellular compartments. GSE115978 contained 7,186 annotated cells and GSE120575 contained 16,291. At least 20 actual cells were required for each specimen and target population before scoring expression aggregates. Patients were the statistical units within each defined treatment and visit stratum. The GSE115978 source Table S1A confirmed that Mel75 and Mel75.1 were repeated material from one patient and that Mel129PA and Mel129PB shared a patient. The two Mel75 specimens were reconstructed and scored separately before equal patient averaging. Patient counts after this aggregation are shown in Table 1.

For anti-PD-1 monotherapy in GSE120575, baseline glycolysis and IFNα ssGSEA scores were positively correlated in CD8 T cell aggregates from 12 patients (Spearman ρ = 0.615, asymptotic P = 0.033, two-test Benjamini-Hochberg P = 0.066). The additional permutation analysis gave an adjusted P value of 0.075. The baseline myeloid correlation was 0.543 in six patients and was treated as descriptive. On-treatment correlations were 0.804 in 19 CD8 T cell patients and 0.773 in 11 myeloid patients (Fig 4). Only five CD8 and two myeloid patients contributed both visits. These separate estimates consequently do not demonstrate an increase caused by treatment or a difference between cell types.

**Fig 4.**
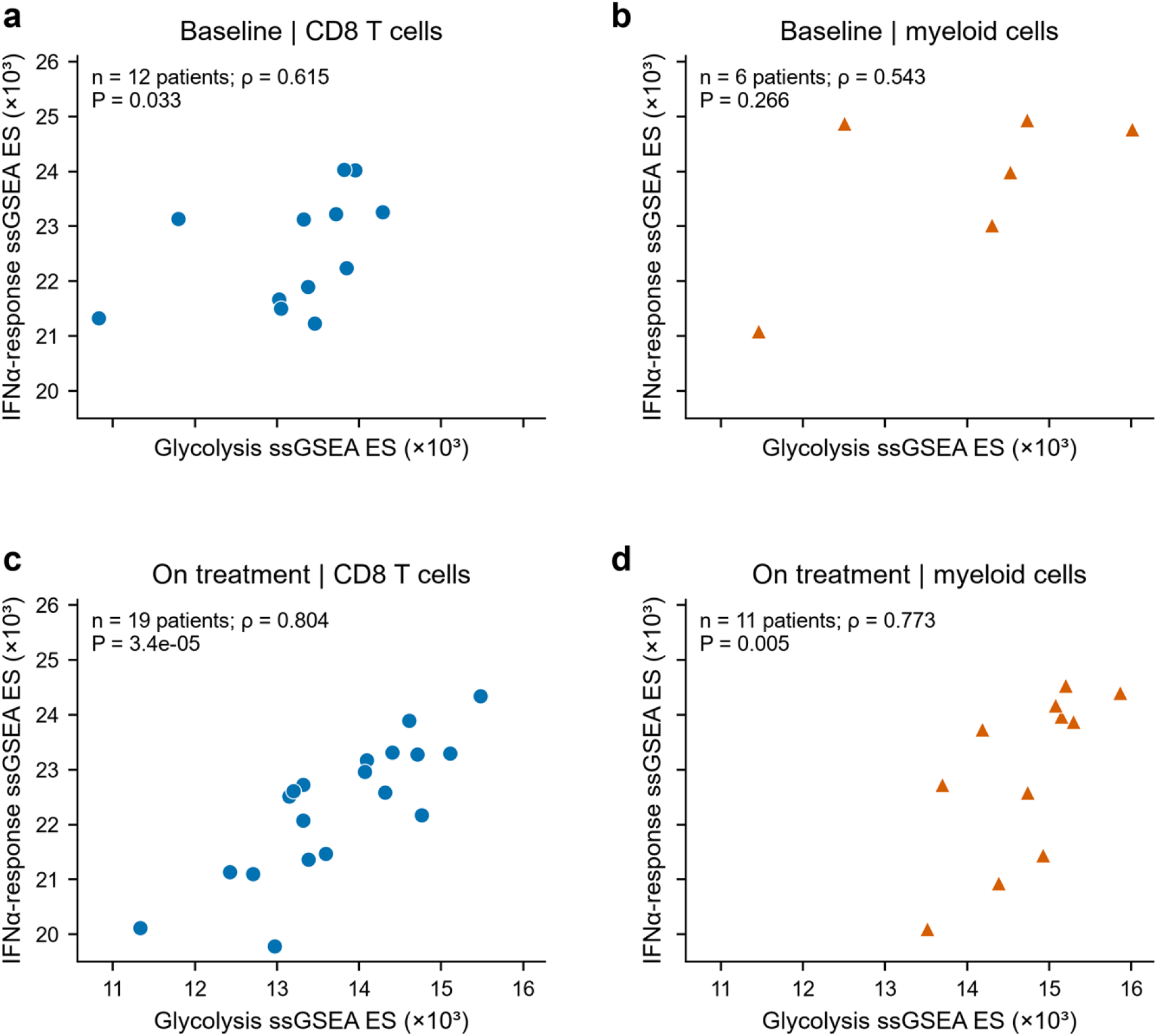
Patient level correlations in GSE120575 immune cells. a,b, Baseline anti-PD-1 samples comprising CD8 T cell aggregates from 12 patients and myeloid aggregates from six patients. c,d, Anti-PD-1 samples collected during treatment from 19 and 11 patients, respectively. Points represent averages of eligible lesion scores for each patient, with a minimum of 20 actual cells for each specimen and target group. Axes show ssGSEA ES scaled by 10^3^ and use common limits. Spearman correlations and unadjusted two-sided P values are annotated. The two baseline comparisons have Benjamini-Hochberg-adjusted P values of 0.066 and 0.266. The small myeloid baseline group is descriptive. Post-treatment analyses are exploratory. The panels compare patients within each time stratum and do not test a change induced by treatment or a difference between cell types. P values for the retained scores shown in this figure use the original asymptotic analysis. Additional permutation results and alternative scoring/backgrounds are presented separately in Fig 5 and Supplementary Table S5.

To assess the dependence on score definition, we calculated an independent score based on mean percentile ranks with average ranks for tied values. Permuting the gene order did not change the numerical results. With the full GSE120575 background, the baseline CD8 correlation was 0.573 (patient bootstrap 95% interval, −0.141 to 0.958). Using the background of 21,957 genes shared by both single-cell datasets reduced the estimate to 0.126 (−0.541 to 0.812). The baseline myeloid estimate changed from 0.543 to 0.029. For on-treatment CD8 and myeloid cells, estimates of 0.746 and 0.809 using the full background changed to 0.489 and 0.436 with the common background (Fig 5, Supplementary Table S5). Changing the background altered both the ranking reference and some gene set coverage. The baseline correlations were therefore sensitive to the ranking background, with substantial uncertainty around both estimates.

**Fig 5.**
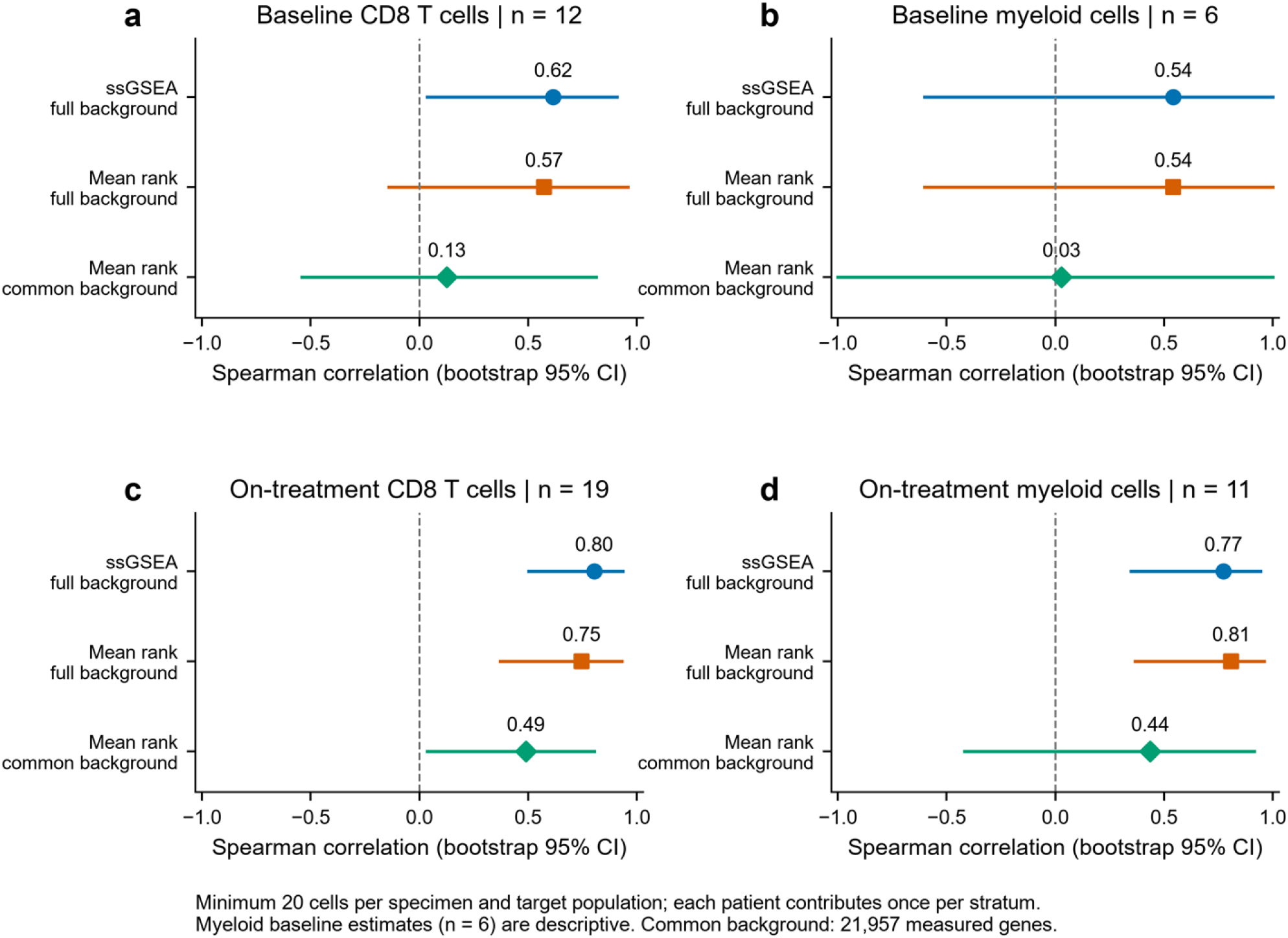
Comparison of single cell scoring methods and gene backgrounds. a to d, Patient-level Spearman correlations in GSE120575 baseline CD8 T cells (12 patients), baseline myeloid cells (six), on-treatment CD8 T cells (19) and on-treatment myeloid cells (11). Dots show ssGSEA, independent mean percentile rank over the full background of each dataset, and independent mean percentile rank over the common background of 21,957 genes. Lines are percentile 95% intervals from 5,000 patient bootstraps. Baseline myeloid estimates are descriptive because of the small patient count. All scores use a minimum of 20 actual cells per specimen and target type. Background changes alter both the rank reference and some gene set coverage. The plot does not test differences between methods or time points. Each sensitivity uses the same underlying patients within its stratum.

GSE115978 provided a further exploratory comparison after correcting patient identities. Among treatment-naive patients, correlations were −0.333 in malignant cells (nine patients), 0.345 in CD8 T cells (ten patients) and −0.400 in myeloid cells (four patients). The previously treated CD8 estimate was 0.291 in 13 patients. Cell thresholds of 10, 20 and 50 and alternative scores changed several estimates, and all eligible results were retained. These small groups differed in treatment history and did not constitute an independent validation of anti-PD-1 efficacy. The cellular correlations varied across scoring settings, and the available patient groups did not identify a consistent cellular source for the clinical association.

## Discussion

The state with high glycolysis and low IFNα response did not show a consistently adverse association with disease control during PD-1 blockade. GSE91061 gave a positive disease control association, GSE78220 gave an estimate in the opposite direction, and the additional Liu estimate was close to one. Neither initial comparison met the Holm-adjusted threshold. Continuous models also provided no consistent evidence of an adverse interaction. The broad confidence intervals remain compatible with clinically relevant effects in either direction. The results do not establish equivalence between the joint state and the other states.

These findings differ from the simple adverse relationship suggested by the mechanism of STAT1 pyruvylation. Zuo and colleagues measured a biochemical modification in experiments involving high glucose and antiviral responses [6]. The present analysis measured gene expression in melanoma biopsies. Pyruvate concentration and STAT1 Lys201 modification were unavailable. Other melanoma studies also show that metabolism and interferon transcription need not change together. Harel and colleagues linked metabolic proteins with antigen presentation and treatment response [1]. Mahmood and colleagues observed increased glycolytic flux after mitochondrial DNA perturbation, while transcriptomes from malignant cells in vivo showed increased interferon programmes together with reduced glycolysis signatures [2]. Chronic tumour interferon signalling can also support adaptive resistance [7]. The current cohort comparisons therefore test a transcriptional hypothesis without validating or refuting the biochemical mechanism.

Previous computational studies have already examined metabolic predictors of checkpoint blockade response, including immune cell states from GSE120575 [3]. We examined whether the specified combination of glycolysis and IFNα transcription was consistently associated with disease control. To interpret this comparison, we examined tumour purity and single-cell expression. IFNα scores correlated with RNA-derived immune scores and with an independent DNA-derived estimate of tumour purity. The latter association is compatible with contributions from nonmalignant cells. Tumour purity does not measure immune cell abundance, and the analysis does not establish why the clinical estimates differed between cohorts. The substantial IFNα/IFNγ gene overlap also prevents a cytokine-specific interpretation.

The longitudinal analysis showed higher on-treatment IFNα transcription among patients with disease control after baseline adjustment. Earlier work also examined pathway signatures during treatment [8]. Both studies considered expression at a defined stage of treatment. The present association does not establish a baseline response predictor. The unadjusted difference in score changes was imprecise, and patients with paired biopsies had lower baseline glycolysis than those without paired profiles. In the single-cell analyses, the positive baseline CD8 correlation weakened when the alternative rank score used a common gene background. The confidence intervals were wide, so this comparison demonstrates sensitivity of the estimates rather than a formal difference between scoring methods.

Several limitations affect generalization. Joint state groups were small, criteria for response assessment differed and the medians defined a relative state within each cohort. The thresholds have not been tested as fixed cutoffs for new patients. The additional Liu evaluation was specified after the initial results were known, although its own inclusion and scoring rules were fixed before outcome associations were inspected. Its biopsies designated by the source as pre-PD1 ranged from 490 days before treatment to the day of initiation, and all four eligibility scenarios were retained because the publication and released matrix disagreed about quality exclusions. Complete patient correspondence across resources was unavailable. Residual confounding, selected paired biopsies and small single-cell strata further restrict interpretation. Without an untreated comparator, these data cannot distinguish a treatment-specific predictive association from a general prognostic association. Further computational studies will require larger cohorts, more precise biopsy timing and measurements from specific cellular compartments or spatial data. Scoring and outcome rules should be fixed before analysis.

## Conclusion

Across three melanoma cohorts, the defined state with high glycolysis and low IFNα response was not consistently associated with poorer disease control during PD-1 blockade. Bulk IFNα transcription was related to DNA-derived tumour purity, and patient-level cellular correlations varied with the scoring method and gene background. The study does not establish a clinical prediction biomarker or a mechanism involving STAT1 pyruvylation.

## Author contributions

Lili Wu performed the formal analysis and investigation, prepared the figures and wrote the original draft. Wei Yang developed the study concept, methodology, reviewed and edited the manuscript.

## Ethics and consent

This study analysed previously published, de-identified public datasets. No participants were recruited and no new samples were collected. Consent and ethical oversight for the original data collections are described in the source publications.

## Data availability

The public transcriptomic datasets are available from NCBI GEO under accessions GSE91061, GSE78220, GSE115978 and GSE120575. The Liu cohort expression and clinical data are available in the supplementary files of Liu et al. [11]. Figure-level source data, clinical mappings and analysis outputs are supplied in S2 Data. Custom code and verification records are supplied in S3 Code, and processed reconstruction inputs are supplied in S4 to S9 Files. The accompanying README describes extraction into a common directory and the analysis entry point.

## Supporting information captions

S1 Appendix. Supporting methods and Supplementary Tables S1 to S6, including subparts. The appendix documents eligibility, label conflicts, regression models, patient influence, cellular scoring and paired-biopsy selection.

**S1 Fig.**
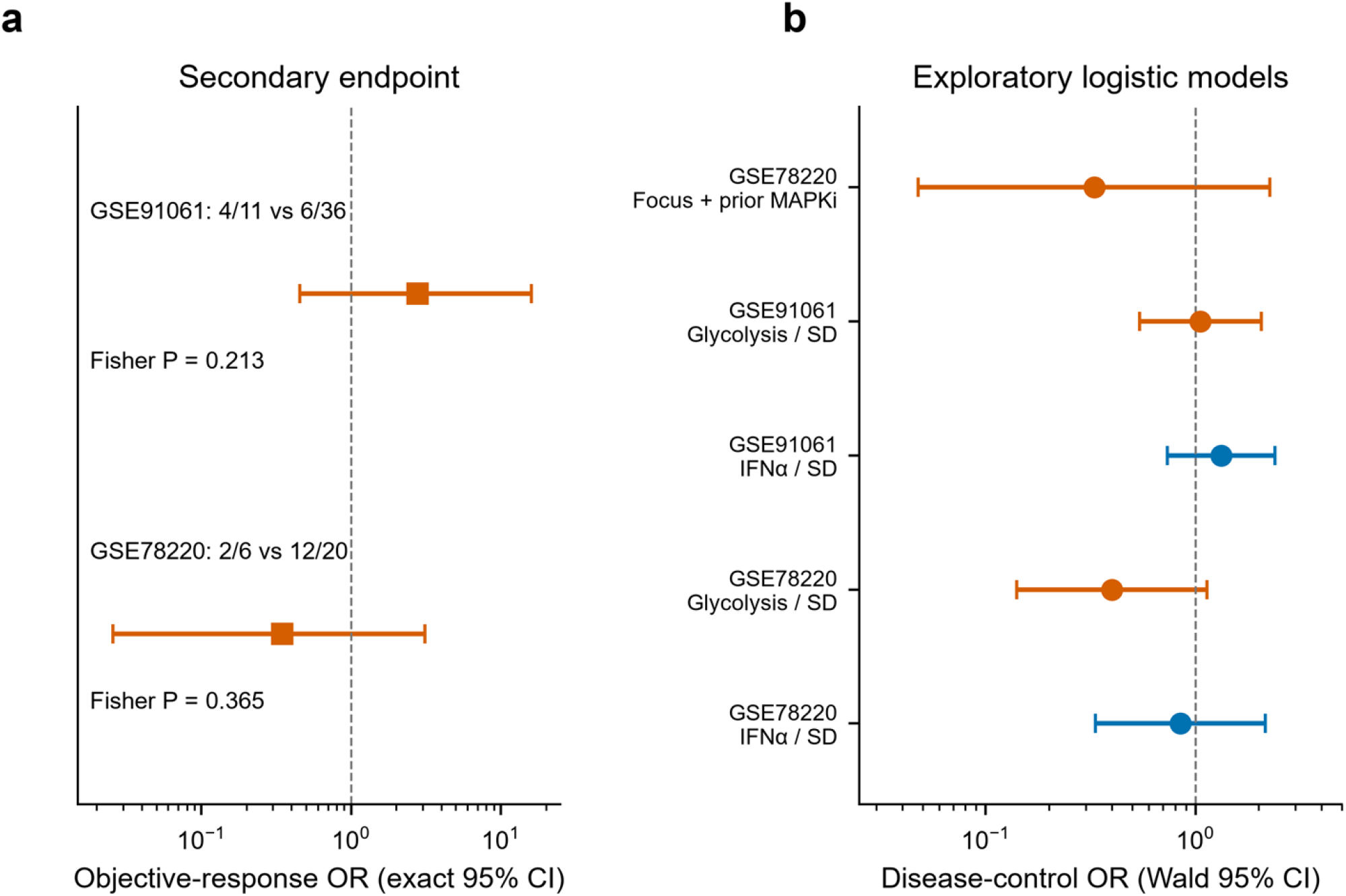
Secondary endpoints and continuous score models. a, Conditional exact odds ratios for objective response (complete or partial response), with exact 95% intervals and two-sided Fisher P values. Labels give response counts in the joint state versus other states. b, Exploratory ordinary maximum likelihood logistic models for disease control. Continuous glycolysis and IFNα coefficients come from a two-score model in each cohort and are expressed per baseline standard deviation. The GSE78220 joint state estimate comes from a separate model adjusted for prior MAPK inhibitor exposure. Bars show Wald 95% intervals. These are not Firth models. All comparisons are exploratory, and intervals are not multiplicity-adjusted.

**S2 Fig.**
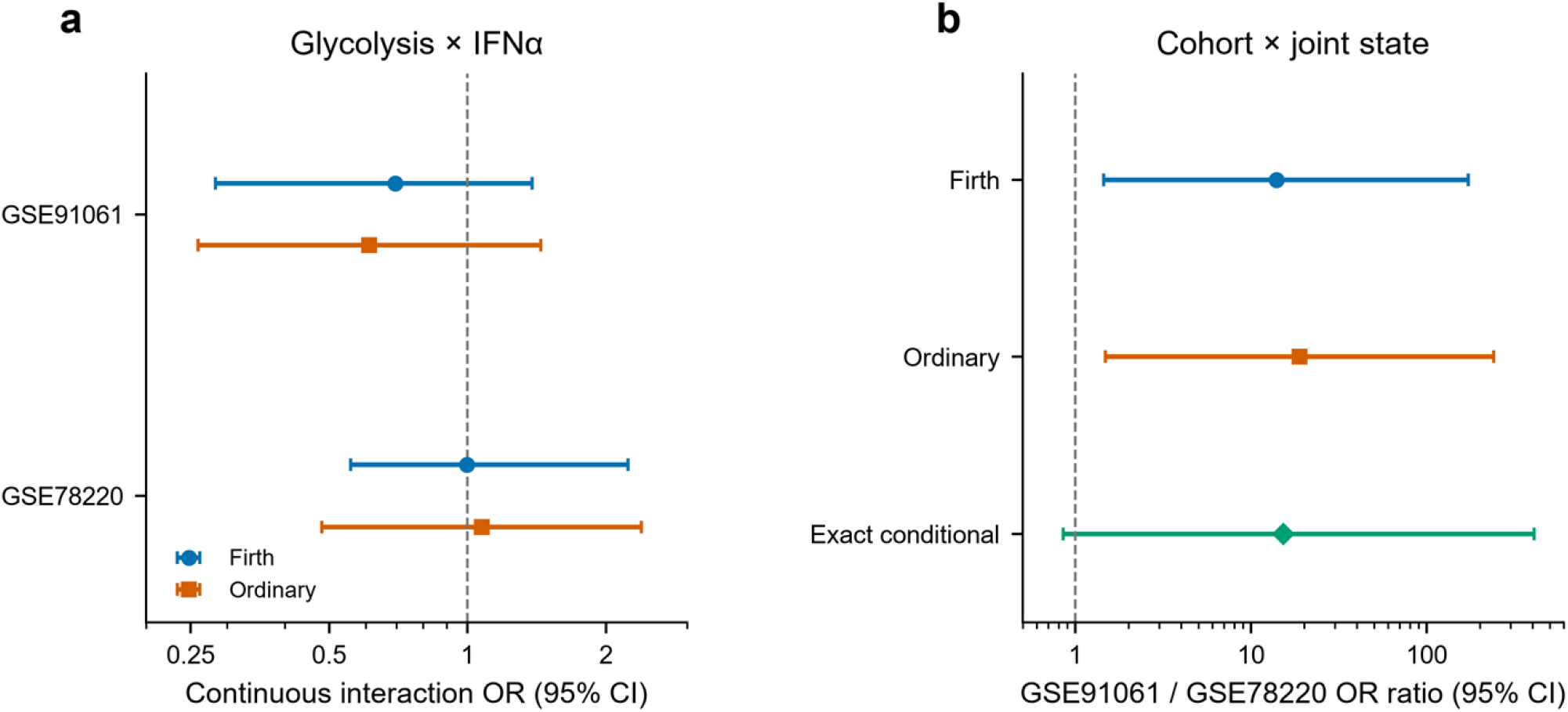
Comparison of interaction estimates. a, Continuous glycolysis × IFNα interaction estimates from the original cohorts. Scores were standardized over all scored baseline patients, and the product was included with both main effects. Firth estimates use penalized profile 95% intervals. Ordinary maximum likelihood estimates use Wald intervals. These are multiplicative interactions on the odds scale, not direct contrasts between the four score groups. b, The ratio of the GSE91061 joint state odds ratio to the GSE78220 joint state odds ratio from an interaction between cohort and state, estimated by Firth, ordinary maximum likelihood and exact conditional enumeration. The Firth raw P value was 0.022 but Holm P was 0.067 across the three added interactions. The exact conditional P value was 0.064. Opposite directions therefore do not establish a precise difference between cohorts that is consistent across methods.

**S3 Fig.**
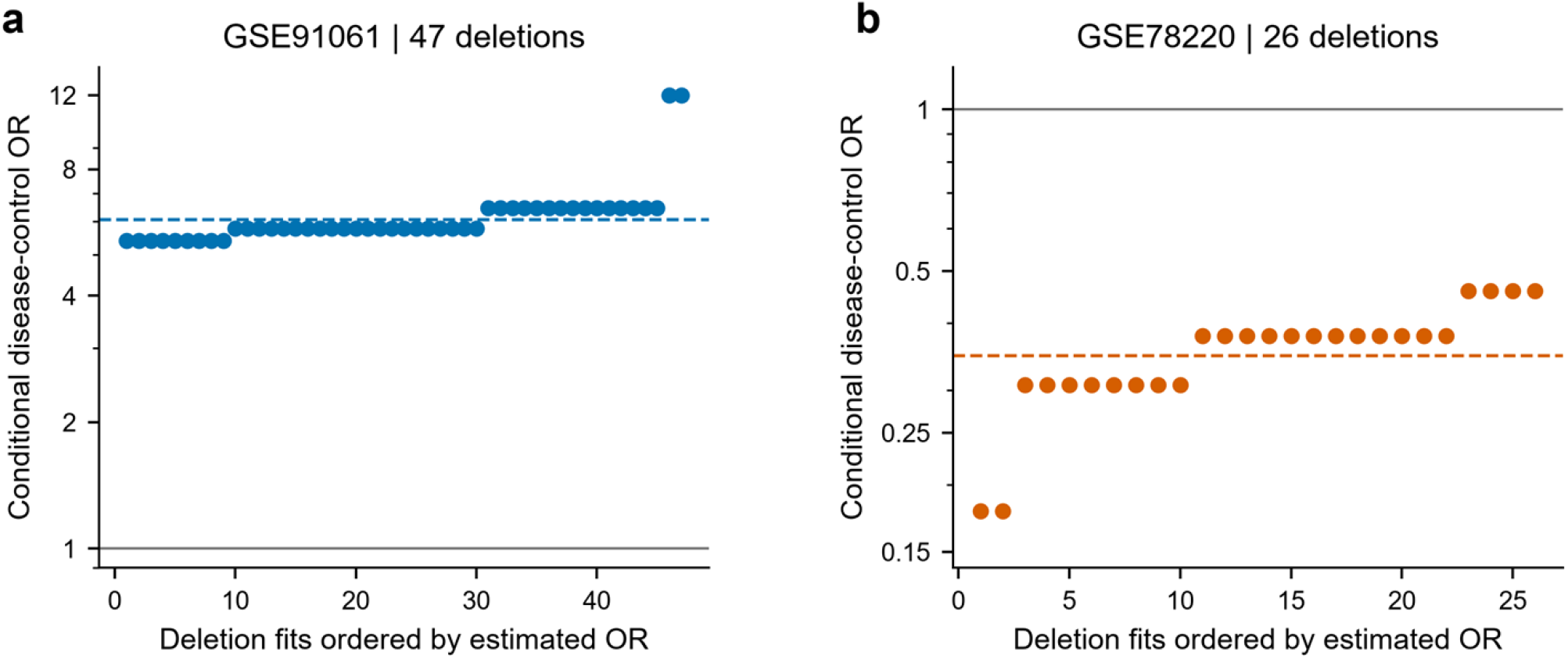
Influence of individual patients on the clinical association. a,b, Conditional disease control odds ratios after deleting each patient once from GSE91061 (47 deletion fits) or GSE78220 (26 fits). Estimates are ordered for display. Dashed coloured lines denote the full-sample conditional odds ratios and grey lines denote OR = 1. Classification thresholds remain fixed at the original all-baseline medians. Deleting a patient does not recalculate the classifier. Neither original effect direction reverses, although confidence and nominal P values change. These are influence analyses, not estimates of cross-validated prediction or clinical utility.

